# How will changes in local climate affect hawksbill hatchling production in Brazil?

**DOI:** 10.1101/410498

**Authors:** Natalie Montero, Maria A.G. dei Marcovaldi, Milagros Lopez–Mendilaharsu, Alexsandro S. Santos, Armando J. B. Santos, Mariana M.P.B. Fuentes

## Abstract

Local climatic conditions can influence sea turtle embryonic development and hatchling viability. Therefore, it is crucial to understand these influences as well as potential ramifications to population stability under future climate change. Here, we examined the influences of five climatic variables (air temperature, accumulated and average precipitation, humidity, solar radiation, and wind speed) at different temporal scales on hawksbill sea turtle (*Eretmochelys imbricata*) hatchling production at ten nesting beaches within two regions of Brazil (five nesting beaches in Rio Grande do Norte and five in Bahia). Air temperature and accumulated precipitation were the main climatic drivers of hawksbill hatching success across Brazil and in Rio Grande do Norte, while air temperature and average precipitation were the main climatic drivers of hatching success at Bahia. Solar radiation was the main climatic driver of emergence rate at both regions. Conservative and extreme climate scenarios show air temperatures are projected to increase, while precipitation projections vary between scenarios and regions throughout the 21^st^ century. We predicted hatching success of undisturbed nests (no recorded depredation or storm-related impacts) will decrease in Brazil by 2100. This study shows the determining effects of different climate variables and their combinations on an important and critically endangered marine species.

## INTRODUCTION

Changes in climate have already impacted the physiology, phenology, behavior, distribution, and reproduction of many species [1-3]. Species that are expected to be most vulnerable are those that are heavily reliant on environmental temperature for their life history traits and/or those that exhibit temperature dependent sex determination (TSD) [4-7]. This is the case for sea turtles, as their life history, physiology, and behavioral traits are heavily influenced by environmental temperature, particularly while their eggs are incubating [8-10]. Successful incubation of sea turtle eggs typically occurs within a specific thermal range of 25°C – 35°C [8], with reduced embryonic development and altered hatchling physiology being observed at extreme temperatures [11-13]. Moisture content also influences embryos with reduced development being observed when conditions are too moist/dry [14-16]. Further, the sex of sea turtle hatchlings is temperature dependent, with warmer temperatures producing a higher proportion of female hatchlings [17, 18]. Thus, any changes to the nesting environment can alter hatchling phenotype and survival, impacting sea turtle populations [19-21].

The impacts of climate change on hatchling production have already been observed through skewed sex ratios, malformations in hatchlings, lowered hatching success, and altered hatchling behavior [13, 22, 23]. Therefore, concern exists over the potential impacts of climate change on hatchling production and population stability [19, 24]. Most studies to date have focused on potential impacts of climate change on the primary sex ratio of hatchlings being produced at nesting grounds [22, 25-27]. However, recent studies have highlighted the fact that the effects of climate change on hatchling production may be of greater concern to population stability [28-30]. Only a few studies to date have investigated the potential impacts of climate change on hatchling production [28, 30, 31], indicating a potential variability in the influence and vulnerability of the different species and populations of sea turtles to various environmental and climatic factors [32]. Indeed, the various species and populations of sea turtles have different thermal tolerances [33, 34] and are also influenced differently by local climate variables [28]. Further, as sea turtle species typically prefer to nest at different locations within a nesting beach (i.e. hawksbills nest on vegetation, whereas loggerheads nest on more open areas) [35-38], it is likely that there is variability on how they can cope with different environmental conditions. This indicates that local climate drivers of hatchling output need to be explored at a species and nesting beach level.

To provide further insights into how sea turtle populations may be impacted by climate change, we expand from previous studies and explore the influences of five different climatic variables (air temperature, accumulated and average precipitation, humidity, solar radiation, and wind speed) on hawksbill sea turtle, *Eretmochelys imbricata*, hatchling production from the Southwest Atlantic Hawksbill Regional Management Unit on the coast of Brazil. This allowed us to determine which variable(s) and combination of variables have the most influence on hatchling production of this critically endangered species, to identify nesting regions that are most susceptible to climate change, and to project future hatching success throughout the 21^st^ century under extreme and conservative climate change scenarios.

## Materials and Methods

### Study site and species

We focused on the hawksbill sea turtle (*Eretmochelys imbricata*) population that nests along the coast of Brazil. This population is part of the Southwest Atlantic Hawksbill Regional Management Unit (RMU) [39]. In Brazil, hawksbills nest at two major nesting regions: southern Rio Grande do Norte (RN) and northern Bahia (BA) [40], which represent the spatial extent of the present study (Fig 1). We used data from five beaches in RN: Cacimbinhas, Chapadao, Madeiro, Minas, and Sibauma and five beaches in BA: Arembepe, Busca Vida, Imbassai, Praia do Forte, and Santa Maria (Fig 1). Although the majority of hawksbill nesting occurs in BA, 42% of hawksbills are hybrids with loggerheads and 2% are hybrids with olive ridley sea turtles [41]. Despite having fewer hawksbill nests, RN has the highest density of nests per kilometer in the South Atlantic Ocean, with some areas experiencing 48.5 nests per kilometer per season [42]. The typical nesting season for hawksbills in RN is November – May, whereas in BA it is October – April [42, 43].

**Fig 1.**
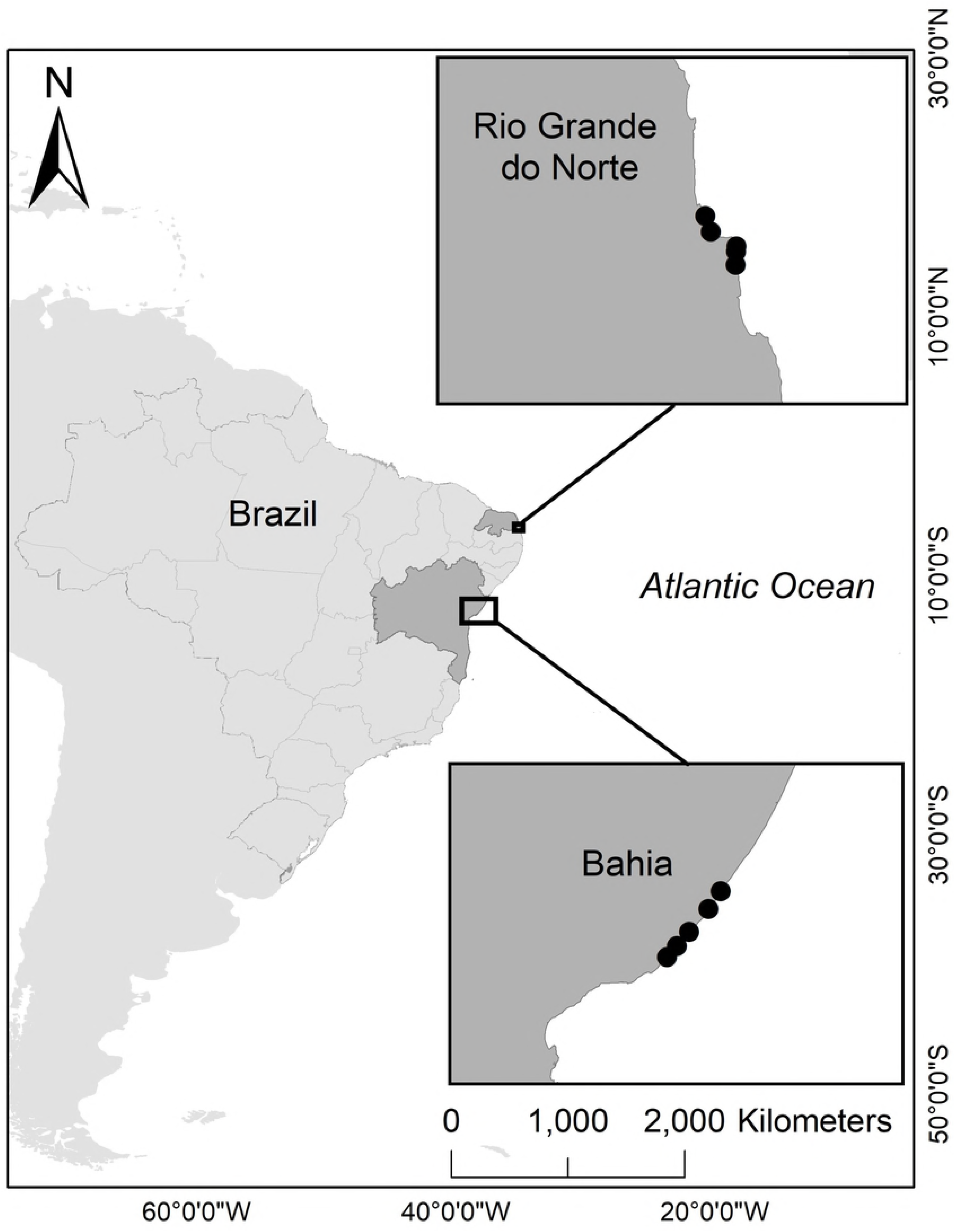
Study sites. Brazilian states (shaded dark gray) considered in this study (north to south): Rio Grande do Norte (RN) and Bahia (BA). Beaches (black points) considered in this study, from north to south: Cacimbinhas, Madeiro, Chapadao, Minas, and Sibauma in RN; and Imassai, Praia do Forte, Arembepe, Santa Maria, and Busca Vida in BA.

### Nest data

Nest data was obtained from Projeto TAMAR, which conducts daily sea turtle patrols in RN and BA during the nesting season. We incorporated nest data from 2005 – 2016 across all 10 nesting beaches listed above. A total of 5,017 hawksbill sea turtle nests were analyzed (RN = 1,334 and BA = 3,683) (Table 1). Variability in the data, for each nesting beach, exists due to logistical and financial constraints involved in monitoring efforts. Only nests that were left *in situ*, had no record of disturbance (i.e. depredation, storm-related impacts, etc.), and had data on location, day laid, hatch date, and hatchling production (hatching success – number of eggs hatched within a clutch- and emergence rate – number of hatchlings that emerged from total hatched eggs within a clutch) were included in this study (Table 1).

**Table 1.** Nest data between and within regions.

| Region | Beach | Number of Nests (Proportions) |
| --- | --- | --- |
| RN | Cacimbinhas | 516 (38.7%) |
| 1,334 (26.6%) | Madeiro | 278 (20.8%) |
|  | Chapadao | 95 (7.1%) |
|  | Minas | 308 (23.1%) |
|  | Sibauma | 137 (10.3%) |
| BA<br>3,683 (73.4%) | Imbassai | 533 (14.5%) |
|  | Praia do Forte | 636 (17.3%) |
|  | Arembepe | 1,414 (38.4%) |
|  | Santa Maria | 587 (15.9%) |
|  | Busca Vida | 513 (13.9%) |

Hawksbill sea turtle nest data from Rio Grande do Norte (RN) and Bahia (BA) considered in this study between 2005 – 2016 (n = 5,017). The number and proportion of nests within each region and across nesting grounds, from north to south.

### Climate data

Local climate data for RN and BA from 2005 – 2016 was obtained from weather stations located in Natal, RN and Salvador, BA, which are maintained by the Brazilian National Institute of Meteorology (INMET; http://www.inmet.gov.br/portal/). These weather stations collect hourly data on air temperature (°C), precipitation (mm), humidity (%), solar radiation (KJ/M2), and wind speed (m/s). Distances between weather stations and nesting beaches are listed in S1 Table. Months where climate data was not available were excluded from any analyses (Table 2).

**Table 2.** Available nest and climate data between regions.

| Rio Grande do Norte months considered<br>(n = 66) | Bahia months considered (n = 67) |
| --- | --- |
| Nov 2005 – Apr 2006; Dec 2006 – May 2007;<br>Nov 2007 – Apr 2008; Nov 2008 – Apr 2009;<br>Nov 2009 – Mar 2010; Nov 2010 – Apr 2011;<br>Dec 2011 – Apr 2012; Nov 2012 – May 2013;<br>Nov 2013 – May 2014; Dec 2014 – Apr 2015;<br>Nov 2015 – May 2016 | Oct 2005 – Dec 2005; Oct 2006 – Mar 2007;<br>Oct 2007 – Jan 2008; Oct 2008 – Apr 2009;<br>Oct 2009 – Apr 2010; Oct 2010 – Apr 2011;<br>Oct 2011 – Apr 2012; Nov 2012 – Apr 2013;<br>Oct 2013 – Mar 2014; Oct 2014 – Apr 2015;<br>Oct 2015 – Apr 2016 |

Projected climate data for RN and BA were obtained from CMIP5 (Coupled Model Intercomparison Project Phase 5) from the KNMI (Royal Netherlands Meteorological Institute) Climate Explorer (https://climexp.knmi.nl/start.cgi), which is a tool used to store climate data and make it easily accessible. We used multi-model means from the extreme RCP (Representative Concentration Pathway) 8.5 scenario and the conservative RCP 4.5 scenario. RCP 8.5 predicts a large increase in global temperatures though the year 2100 due to increasing greenhouse gas concentrations. RCP 4.5 predicts a milder increase in global temperatures through the year 2100 due to predicted future stabilization of greenhouse gas concentrations.

Nest and climate data availability for each region between 2005 – 2016 included in our analyses. The typical nesting season in RN occurs between November – May, while the typical nesting season in BA occurs between October – April.

## Analysis

A one-way ANOVA was used to compare hatchling production between regions and across nesting beaches within each region. Levene’s test, with center mean, found the variances for both hatching success and emergence rate between nesting beaches in RN (p < 0.01, both hatching success and emergence rate) and BA (p < 0.01, both hatching success and emergence rate) to not be homogenous. Therefore, Tamhane’s T2 test was conducted to analyze how hatchling production may differ between nesting beaches within each region.

Generalized Linear Mixed-Effects Models, with the binomial family in package lme4, were used to test the influences of local climate on hatching success and emergence rate. Year was set as the random effect to allow for analysis and predictions across and within years. To best fit these models, hatching success and emergence rate data were presented as successes and failures. For instance, hatching success consisted of the number of eggs that hatched versus those that did not, while emergence rate consisted of the number of hatchlings that emerged from hatched eggs versus those that did not. Corrected Akaike Information Criterion (AICc) was used to identify the best model within each region. Using the best models, hatching success was projected into the future under conservative and extreme climate change scenarios for each region. Emergence rate was not projected due to its potential to be heavily influenced by sand compaction, substrate type, and hatchling performance, which were not measured here [44]. R version 3.4.2 was used for all analyses.

The predictor variables used were: average air temperature (temp), accumulated precipitation (acc.rain), average precipitation (avg.rain), average humidity (humid), average solar radiation (rad), and average wind speed (wind). These predictors were explored at various temporal scales: the month nests were laid (0.climate variable), the month nests were laid and one-month prior (0.1.climate variable), the month nests were laid and two months prior (0.2.climate variable), two months prior to nesting (2.climate variable), and during the incubation period (inc.climate variable). Existent research has shown that air temperature and precipitation as well as humidity have significant influences on hatchling output due to the variability that may exist in moisture throughout the nesting season (i.e. dry and wet seasons) and across regions [15, 28, 38, 45]. Therefore, accumulated precipitation, average precipitation, and humidity predictors were combined with average air temperature during incubation to explore their combined effects on hatchling production. Precipitation projections were presented as mm/day. Therefore, to project accumulated precipitation, these values were converted into mm/month.

## Results

### Hatchling production

Hatching success of undisturbed nests laid between 2005 – 2016 varied across years, regions and nesting beaches (Table 3). BA had the highest average hatching success (76.3% ± 21.6 SD), while the average at RN was 75.2% ± 22.5 SD (Table 3). No significant difference was found in hatching success across years between states (ANOVA, F = 2.8, DF = 1, p = 0.09). A significant difference was found in hatching success across years between nesting beaches in RN (ANOVA, F = 12, DF = 4, p < 0.01) and between nesting beaches in BA (ANOVA, F = 291.2, DF = 4, p < 0.01) (S2 Table). In RN, the beach with the highest hatching success was Madeiro (80% ± 19.3 SD), while Chapadao had the lowest (65.4% ± 27.3 SD) (Table 3). In BA, the beach with the highest hatching success was Arembepe (88.2% ± 12.6 SD), while Praia do Forte had the lowest rates (63.4% ± 23.9 SD) (Table 3). April had the highest average hatching success across both RN (82.3%) and BA (82.6%; Fig 2A). November had the lowest hatching success in RN (72.2%), while October had the lowest average hatching success in BA (69.9%; Fig 2A).

**Table 3.** Hatchling production between and within regions.

| Region | Nesting Beach | Hatching Success (%) |  | Emergence Rate (%) |  |
| --- | --- | --- | --- | --- | --- |
| RN | Cacimbinhas | 75.2% ± 22.5 SD | 77.2% ± 20.3 SD | 88% ± 15.4 SD | 89.2% ± 15.4 SD |
|  | Madeiro |  | 80% ± 19.3 SD |  | 91.2% ± 11.7 SD |
|  | Chapadao |  | 65.4% ± 27.3 SD |  | 81.9% ± 18.5 SD |
|  | Minas |  | 70.7% ± 24.7 SD |  | 84.8% ± 16 SD |
|  | Sibauma |  | 74.6% ± 24.2 SD |  | 88.2% ± 16.4 SD |
| BA | Imbassai | 76.3% ± 21.6 SD | 70.2% ± 24.6 SD | 86% ± 14.1 SD | 83.2% ± 16.9 SD |
|  | Praia do Forte |  | 63.4% ± 23.9 SD |  | 83.2% ± 16.7 SD |
|  | Arembepe |  | 88.2% ± 12.6 SD |  | 90.9% ± 9 SD |
|  | Santa Maria |  | 69.2% ± 21.8 SD |  | 83.2% ± 14.5 SD |
|  | Busca Vida |  | 73.9% ± 18.2 SD |  | 82% ± 14.3 SD |

**Fig 2.**
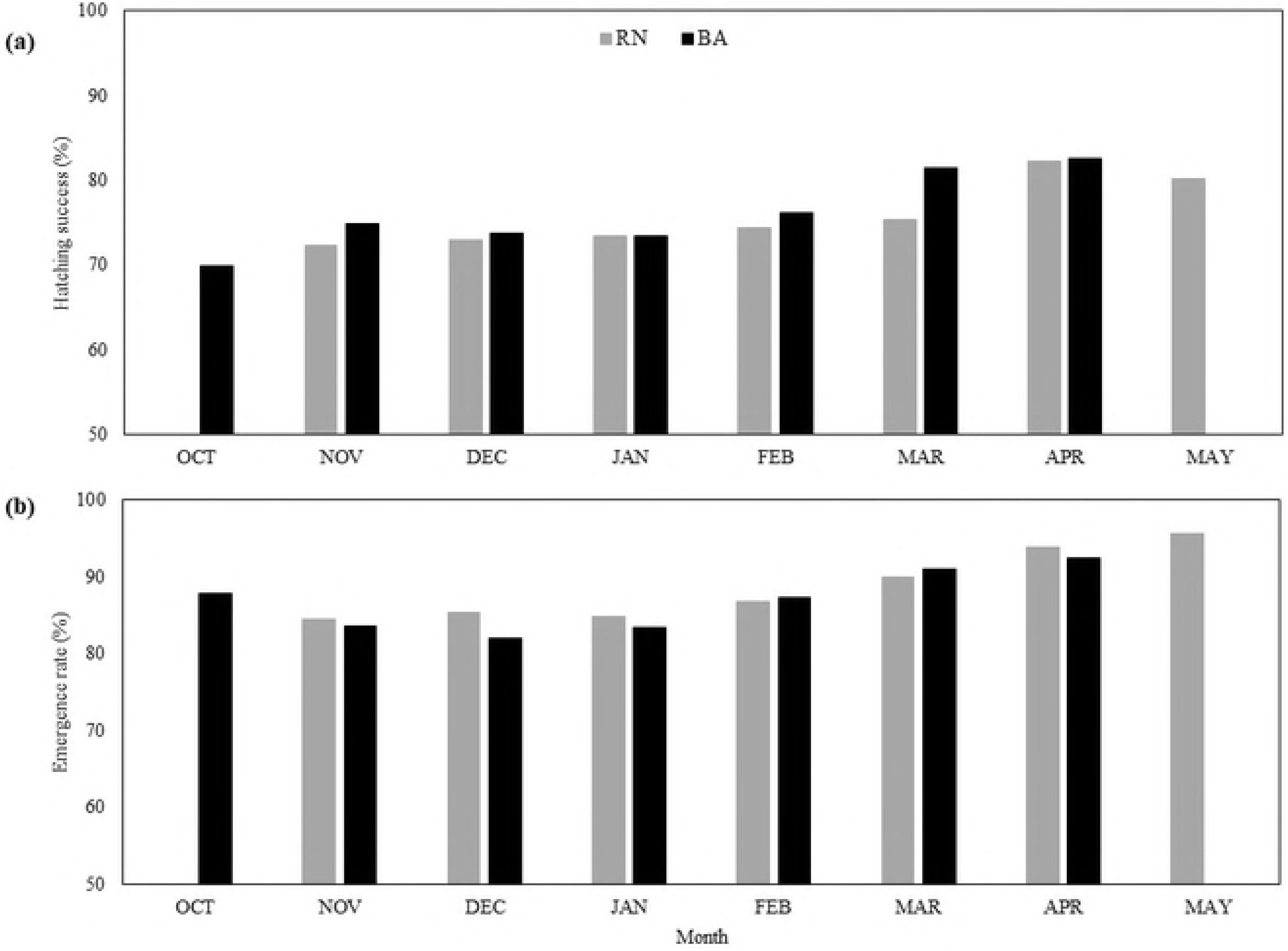
Hatchling production within regions. Monthly averages (bars) for hatching success (A) and emergence rate (B) at RN (light gray) and BA (black), from 2005-2016. The typical nesting season for hawksbills in RN is from November – May, while in BA it is October – April.

Emergence rate of undisturbed nests laid between 2005 – 2016 also varied across years, states, and nesting beaches (Table 3). RN had the highest average emergence rate at 88% ± 15.4 SD, while the average at BA was 86% ± 14.1 SD (Table 3). A significant difference was found in emergence rate across years between states (ANOVA, F = 57.3, DF = 1, p < 0.01) and across years between nesting beaches in RN (ANOVA, F = 13.8, DF = 4, p < 0.01) and between nesting beaches in BA (ANOVA, F = 65.7, DF = 4, p < 0.01) (S2 Table). Similar to hatching success, Madeiro had the highest average emergence rate in RN (91.2% ± 11.7 SD) and Chapadao had the lowest (81.9% ± 18.5 SD) (Table 3). In BA, Arembepe had the highest average emergence rate (90.9% ± 9 SD), while Busca Vida had the lowest (82% ± 14.3 SD) (Table 3). May had the highest emergence rate in RN (95.6%), while April had the highest emergence rate in BA (92.4%; Fig 2B). January had the lowest emergence rate in RN (84.8%), while December had the lowest emergence rate in BA (82.1%; Fig 2B).

Average hatching success and emergence rate from 2005 – 2016, with their respective standard deviations, at each nesting beach in Rio Grande do Norte (RN) and Bahia (BA), from north to south, throughout the study period.

### Local climate

Hawksbill sea turtles nest during the warmer months of the year throughout RN and BA (October – May) (Fig 3). RN is the warmest of the two regions being closer to the equator than BA (Fig 3a). In RN, the warmest month is February (27.64°C ± 0.4), while in BA the warmest month is March (27.35°C ± 0.64) (Fig 3A). During February and March, adult females are still nesting in high proportions, but hatchlings are incubating and emerging from nests (Fig 3A). The wettest month in both RN and BA is May, with an accumulated precipitation of 200.92 mm ± 153 in RN and 190.35 mm ± 129 in BA (Fig 3B). Likewise, May had the highest average precipitation in both RN and BA, with 0.31 mm/day in RN and 0.26 mm/day in BA (Fig 3C). In May, there is no nesting in BA and nesting is finishing in RN, but hatchlings are still incubating and emerging (Fig 3C). RN has more solar radiation than BA, likely due to its proximity to the equator (Fig 3D). The month with the highest average solar radiation in RN is October (1907.7 KJ/M^2^ ± 198.7), while in BA it is January (1754.2 KJ/M^2^ ± 158.2) (Fig 3D). In RN, there is no nesting in October, when solar radiation is highest (Fig 3d). On the other hand, January is when solar radiation and nest proportions are at their highest in BA (Fig 3D). BA is more humid than RN and May is the most humid month in both RN (79.2% ± 2.05) and BA (80.3% ± 5) (Fig 3E). In May, nesting has ended in BA and is ending in RN, but hatchlings are still incubating and emerging (Fig 3E). RN is windier than BA, with October being the windiest month in both RN (5.29 m/s ± 0.5) and BA (1.82 m/s ± 0.34) (Fig 3F). There is no nesting in October in RN, while October has the lowest proportion of nests in BA (Fig 3F).

**Fig 3.**
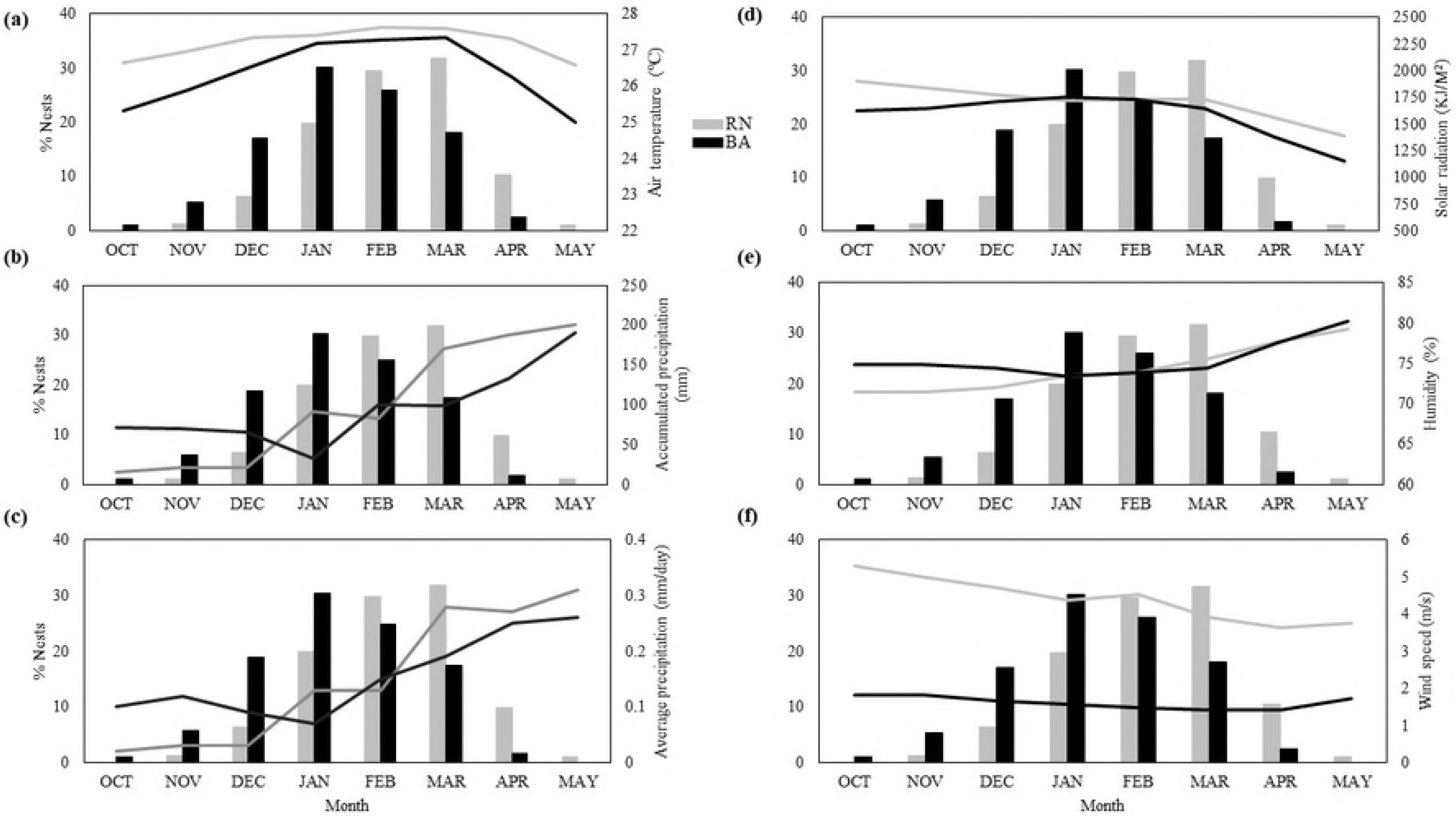
Climatic conditions within regions. Monthly climatic conditions (lines) and proportion of nests laid (bars) within RN (light gray) and BA (black) for each month within the nesting season (October – May) between 2005 – 2016. Climate variables are listed as follows: (A) air temperature, (B) accumulated precipitation, (C) average precipitation, (D) humidity, (E) solar radiation, and (F) wind speed. The typical nesting season for hawksbills in RN occurs between November – May, while the typical nesting season in BA occurs between October – April.

### Effects of local climate on hatchling production output

Throughout Brazil, the model with the lowest AICc and high significance for hawksbill hatching success was average air temperature during incubation in combination with accumulated precipitation during the month nests were laid and two months prior (p < 0.001 for both parameters; Fig 4A, S3 Table). This model showed lower hatching success when conditions were warm and dry (Fig 4A). For emergence rate throughout Brazil, the model with the lowest AICc and high significance was average solar radiation during incubation (p < 0.001; Fig 4B, S3 Table). Emergence rate decreased with increasing solar radiation (Fig 4B).

**Fig 4.**
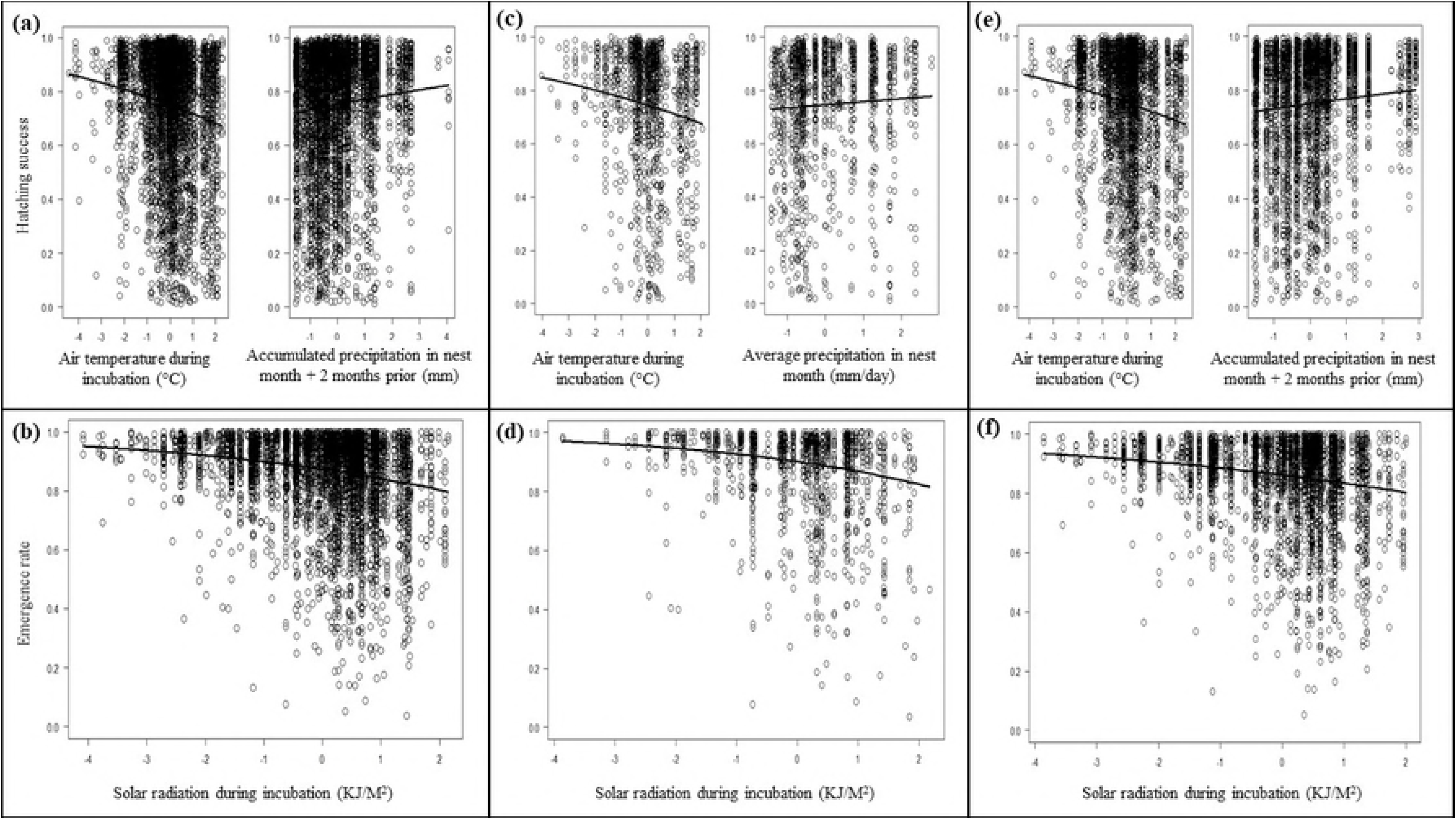
Model results. Best fit models describing hatching success (A, C, E) and emergence rate (B, D, F) across Brazil (A, B) and within RN (C, D) and BA (E, F).

Across RN, the model with the lowest AICc and high significance for hatching success was average air temperature during incubation in combination with average precipitation during the month nests were laid (p < 0.001 for both parameters; Fig 4C, S3 Table). This model indicated that warmer and drier conditions decreased hatching success (Fig 4C). The model with the lowest AICc and high significance for emergence rate across RN was average solar radiation during incubation (p < 0.001) (Fig 4D, S3 Table). Here, higher solar radiation decreased emergence rate (Fig 4D).

The model with the lowest AICc and high significance for hatching success across BA was average air temperature during incubation in combination with accumulated precipitation during the month nests are laid and 2 months prior (p < 0.001 for both parameters; Fig 4E, S3 Table). This model indicated that warmer and drier conditions decreased hatching success (Fig 4E). For emergence rate across BA, the model with the lowest AICc and high significance was average solar radiation during incubation (p < 0.001; Fig 4F, S3 Table). Here, higher solar radiation decreased emergence rate (Fig 4F).

### Climate projections

By 2100, air temperatures at RN are projected to increase throughout the nesting season by 1.4 – 4°C with November being the warmest month under RCP4.5 at 29.4°C and 31.5°C under RCP8.5 (Fig 5A). Average precipitation projections vary throughout the 21^st^ century, however there is a general increase of 0.11 – 1.86 mm/day throughout the nesting season in RN (Fig 5B). November is projected to be the wettest month by 2100 under RCP4.5 at 1.03 mm/day and under RCP8.5 at 1.89 mm/day (Fig 5B). Air temperatures in BA are projected to increase throughout the nesting season by 1.8 – 5°C with March being the warmest month under RCP4.5 at 29.3°C, but the warmest month under RCP8.5 is projected to be January at 32°C (Fig 5C). Accumulated precipitation is projected to vary, but there is a general increase of 0.9 – 22.3 mm throughout the nesting season in BA (Fig 5D). May is projected to be the wettest month by 2100 under RCP4.5 at 185.7 mm and under RCP8.5 at 180.1 mm (Fig 5D).

**Fig 5.**
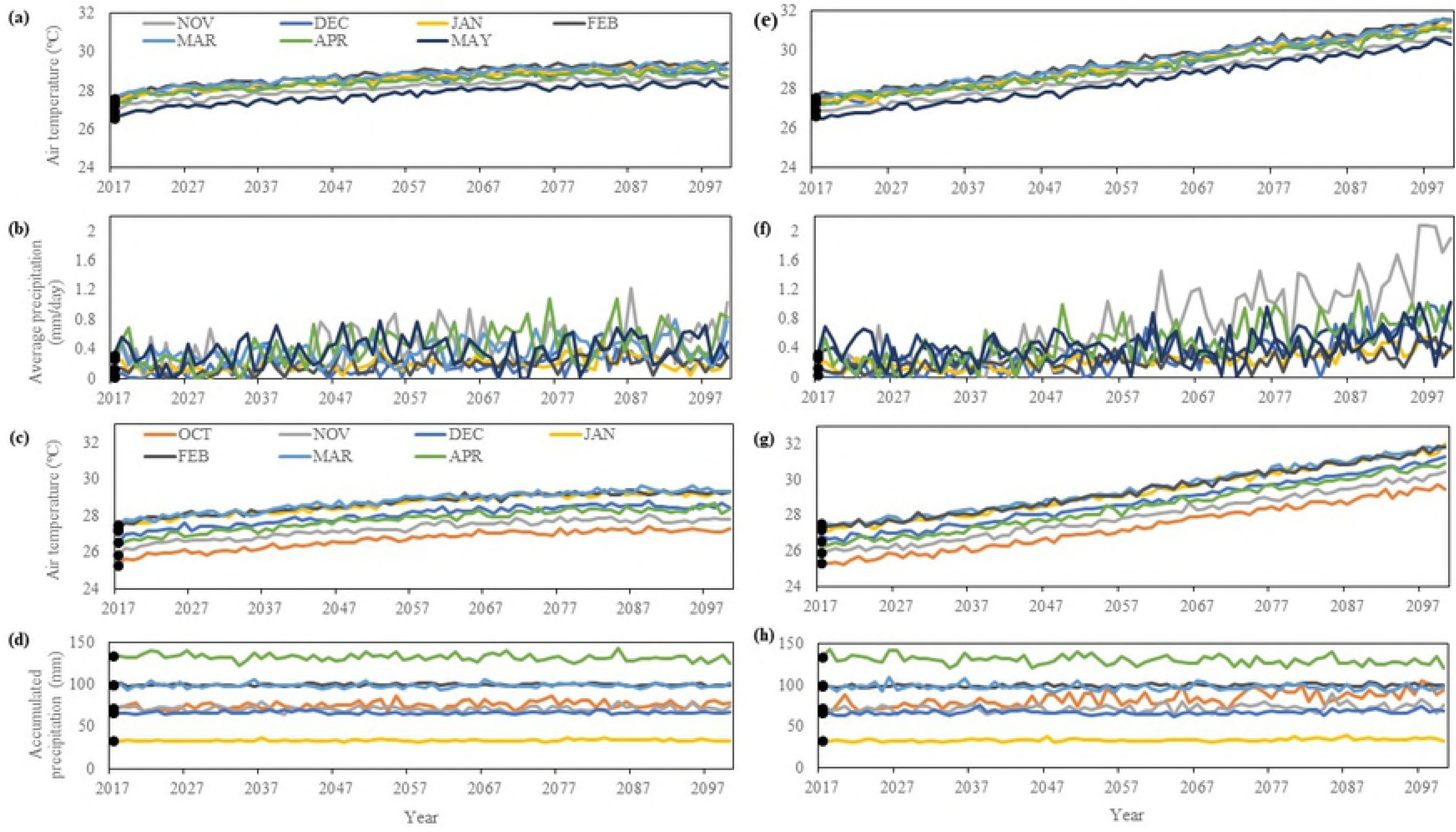
Climate projections. Historic average (black points) and projected values (lines) for each month in the nesting season for air temperature (A, E) and average precipitation (B, F) at RN and air temperature (C, G) and accumulated precipitation (D, H) BA under the conservative RCP4.5 scenario (left panels) and the extreme RCP8.5 scenario (right panels). The typical nesting season for hawksbills in RN occurs between November – May, while in BA it is October – April.

### Hatching success projections

We used the best fit models describing hatching success at RN (air temperature during incubation in combination with average precipitation during the month nests were laid) and BA (air temperature during incubation in combination with accumulated precipitation during the month nests were laid and two months prior) as well as our climate deltas to project hatching success throughout the 21^st^ century. Our models predicted average hatching success to decrease in both RN and BA thought the 21^st^ century. By 2100, hatching success in RN is projected to decrease from an average across the study period of 75.2% to 72.6% under RCP4.5 and to 70.9% under RCP8.5 (Fig 6A). Hatching success in BA is projected to decrease from an average across the study period of 76.3% to 65.1% under RCP4.5 and 69.6% under RCP8.5 (Fig 6B).

**Fig 6.**
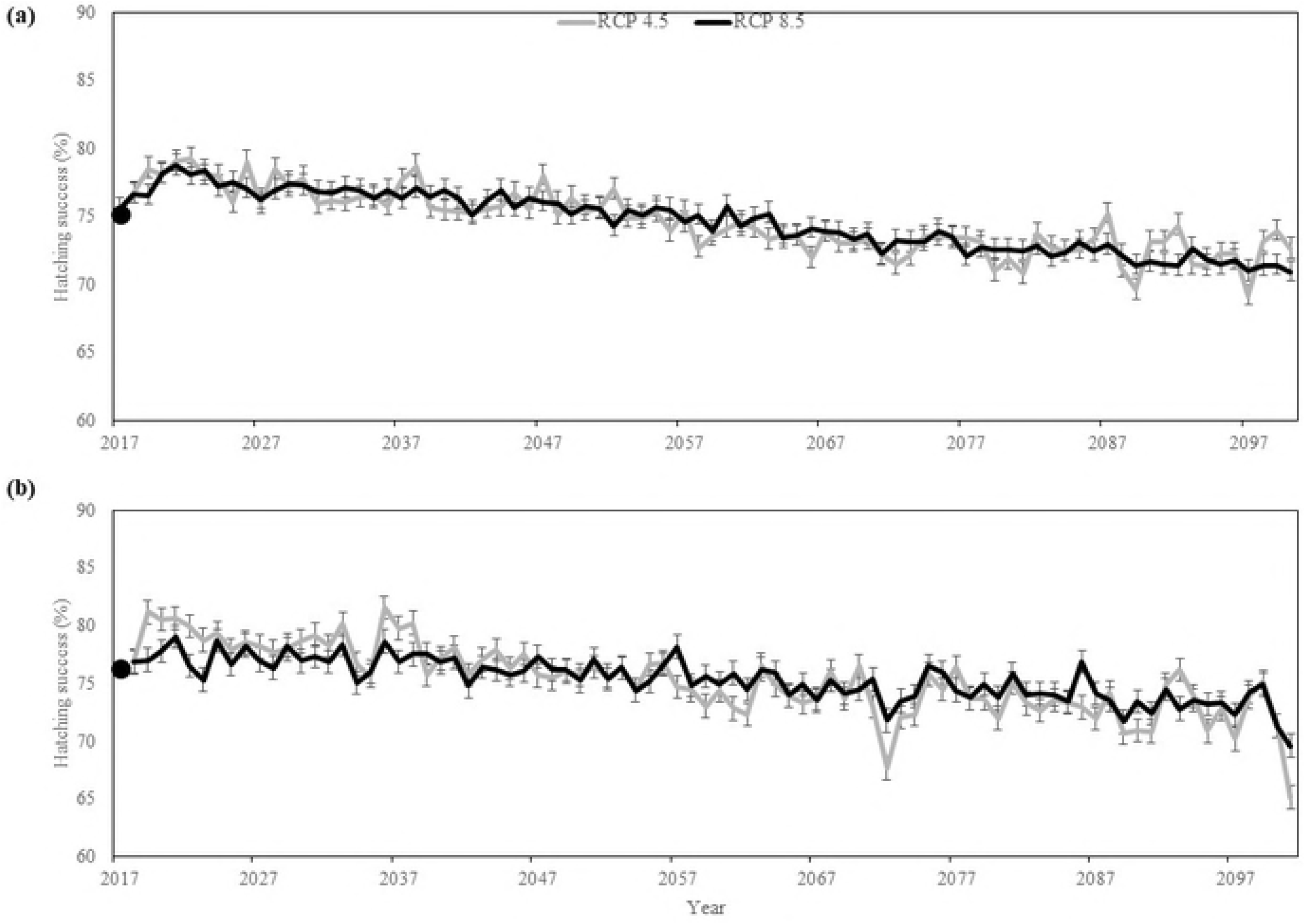
Hatching success projections. Historic average (2005 – 2016, black points) and projected values (lines) for hatching success in RN (A) and BA (B) under the conservative RCP4.5 scenario (light gray) and extreme RCP8.5 scenario (black). Bars represent confidence intervals.

## Discussion

Hawksbill nesting beaches in Brazil are projected to experience decreases in hatching success of up to 11% by 2100 due to warming air temperatures and increases in precipitation. Air temperature in combination with accumulated precipitation were found to be the main climatic drivers of hawksbill hatching success across Rio Grande do Norte, while air temperature in combination with average precipitation were the main climatic drivers of hatching success in Bahia. Across all regions, higher hatching success was observed in cooler and wetter conditions. A positive influence of precipitation on hatching success was found across Brazil and within both regions, which may be a reflection of an absence of adequate moisture levels at these nesting grounds, where most of the nesting (January to March) occurs during the drier months of the season [38, 45]. Consequently, projections show a lower decrease in hatching success under the more extreme climate change scenario, RCP8.5, as it projects higher increases in precipitation when compared to the conservative scenario (RCP4.5). Wetter, more moist conditions may offset increases in temperature maintaining cooler sand temperatures [15, 46-48]. However, as climate change progresses and conditions become wetter, there is a potential for further reductions in hatchling production, since very wet conditions can result in soil saturation or a rise of the water table level displacing air between sand particles, suffocating embryos and resulting in clutch failure [28, 49, 50]. This is the case for loggerhead turtles in Brazil, where a negative effect of precipitation on loggerhead hatching success was found for nesting beaches in Bahia that are shared between loggerhead and hawksbills, specifically in Praia do Forte and Santa Maria [51]. This is likely a reflection of the fact that the peak for loggerhead turtle nesting in Brazil overlaps with the wettest months (October to December) of the season and these areas already experience high levels of rain [38, 45, 51]. Interestingly, although hawksbill and loggerhead turtles have some common nesting areas in Bahia, Brazil [43, 52], the relationship between precipitation and hatching success for these species are different. This variability may be driven by differences in nesting seasonality and nest placement between the species.

Hawksbill turtles are known to nest near or within vegetation, further from the high-water mark than loggerhead turtles [35-37, 53]. Sand temperatures tend to be significantly cooler than in open areas [54, 55]. Indeed, mean incubation duration for hawksbill turtles is longer than for loggerhead turtles in Bahia, Brazil, indicating that hawksbill eggs are incubating at cooler temperatures [43, 52]. Sea turtle nesting behavior and historical nest placement may drive the tolerance of species to thresholds of incubating parameters (i.e. moisture, temperature) and drive the adaptive differentiation of species at fine spatial scales [56]. Indeed, it was found that hatchlings of females nesting on a naturally hot (black sand) beach survived better and grew larger at hot incubation temperatures compared to the offspring of females nesting on a cooler (pale sand) beach nearby [56]. Thus, it could be speculated that the thermal tolerances of loggerhead eggs incubating in Brazil are higher than those for hawksbill turtles. Furthermore, there is a high prevalence of hybridization between hawksbill and loggerhead sea turtles with 42% being morphologically assigned as hawksbills [41]. Hatching success between hybrids and their parental species are not significantly different, however hybrid emergence rates were found to be lower than the parental species [57]. Therefore, these hybrids may endure in Brazil however, their responses to climate remain unknown.

With projected changes in climate, it is important to understand the climatic thresholds for various species of sea turtles to better project potential impacts and to inform management. Thermal tolerances are known to vary between species of sea turtles [32] and are also likely to vary between populations and within nesting grounds as a reflection of strong site fidelity by individual females. Future studies should also explore whether sea turtles are able to locally adapt to other environmental conditions (i.e. moisture) and integrate these findings into models that predict population responses to climate change. Similarly, a better understanding of the interacting and synergetic effects of various environmental conditions (temperature, moisture, etc.) is necessary. The effects of temperature on hatchling output is relatively well understood [58-60] with recent advancements in our understanding of the effects of moisture [15, 48, 61]. However, other less studied environmental variables may be affecting the reproductive output of sea turtles at nesting grounds. Here, we also explored how solar radiation and wind speeds may influence hatchling output. Although our study suggests air temperature and precipitation to be most significant, solar radiation, was a good predictor and had a negative effect on emergence rate across Brazil and within both regions. High solar radiation can enhance the effect of warm air temperatures by warming the sand, and thus heating nests beyond the thermal threshold, potentially negatively impacting hatchling production [62]. Further, sand temperatures that are too warm, and consequently very dry, can reduce successful hatchling emergence since it can cause nests to cave-in making it difficult for hatchlings to emerge [12, 63]. Ultimately, if nesting and incubating conditions are not favorable, as a consequence of climate change or other threats, sea turtles will need to respond and adapt accordingly.

Sea turtles have been around for millions of years and have persisted through dramatic changes in past climates, demonstrating their ability to adapt to changing conditions [20, 64], by: 1) changing the distribution of their nesting grounds, nest site choice, and nest depth; 2) adapting *in situ* by adjusting their pivotal temperature; and 3) shifting their nesting to cooler months of the year [21, 65-70]. It is suggested that range shifts may offer one of the most promising avenues for adaptation in marine turtles [62, 64] since earlier nesting and changes in nest-site choice can quickly offset and counteract projected impacts to species with temperature-dependent sex determination [4, 71, 72]. Range shifts have already been observed for leatherback and loggerhead turtles as response to warming temperatures [69, 73, 74].

Unfortunately, the future adaptive capacity of sea turtles may be hindered by the reduction of available nesting grounds from rises in sea-levels and the ever-expanding development along coasts [75-77]. Thus, management efforts should focus on ensuring that nesting grounds are available to increase the resilience of sea turtle populations to climate change [78, 79]. This can be achieved by protecting nesting beaches (i.e. conservation easements, reducing coastal development, enforcing wildlife laws and regulations), limiting and regulating development (i.e. implementing setback regulations), and by maintaining the nesting habitat (i.e. maintain native vegetation, minimize the use of shoreline hardening structures) [54, 77, 79-81].

## Acknowledgements

We would like to thank the volunteers, interns, and staff at Projeto TAMAR and INMET who contributed to the collection and distribution of the nest and climate data used in this study.

## Supporting Information Captions

**S1 Table. Weather station locations.** Distances of INMET weather stations from nesting grounds in each region considered in this study, from north to south.

**S2 Table. Statistical differences in hatchling production between and within regions.** Results of Tamhane’s T2 test for statistical differences in hatching success and emergence rate between nesting beaches within Rio Grande do Norte (RN) and Bahia (BA), Brazil, from north to south. Statistically significant p – values are indicated in bold.

**S3 Table. Statistical results of climate influences on hatchling production.** Results of Generalized Linear Mixed-Effects Models for local climate influences on hatching success (HS) and emergence rate (ER) across Brazil as well as within Rio Grande do Norte (RN) and Bahia (BA). For these models, the binomial family was specified, and the year nests were laid was the random effect. Model parameters included: average air temperature (temp), accumulated precipitation (acc.rain), average precipitation (avg.rain), average humidity (humid), average solar radiation (rad) and average wind speed (wind). The temporal scales used were: the month nests were laid (0.climate variable), the month nests were laid and one-month prior (0.1.climate variable), the month nests were laid and two months prior (0.2.climate variable), two months prior to nesting (2.climate variable), and during the incubation period (inc.climate variable). The models with the lowest AICc values and high significance are highlighted in gray. P – values for combined models are presented for each parameter in the order the model is written.

